# No need for extensive artifact rejection for ICA - A multi-study evaluation on stationary and mobile EEG datasets

**DOI:** 10.1101/2022.09.13.507772

**Authors:** M. Klug, T. Berg, K. Gramann

**Affiliations:** Young Investigator Group Intuitive XR, Neuroadaptive Human-Computer Interaction, Institute of Medical Technology, BTU Cottbus-Senftenberg, Cottbus, Germany; Biopsychology and Neuroergonomics, Institute of Psychology and Ergonomics, TU Berlin, Berlin, Germany

**Author notes:** authors contributed equally (M. Klug, T. Berg).

**Keywords:** electroencephalography, mobile brain/body imaging, signal processing, artifact removal, preprocessing, independent component analysis

## Abstract

**Objective:** Electroencephalography (EEG) studies increasingly make use of more ecologically valid experimental protocols involving mobile participants who actively engage with their environment (MoBI; Gramann et al., 2011). These mobile paradigms lead to increased artifacts in the recorded data that are often treated using Independent Component Analysis (ICA). When analyzing EEG data, especially in a mobile context, removing samples regarded as artifactual is a common approach before computing ICA. Automatic tools for this exist, such as the automatic sample rejection of the AMICA algorithm (Palmer et al., 2011), but the impact of the two factors movement intensity and the automatic sample rejection has not been systematically evaluated yet.

**Approach:** We computed AMICA decompositions on eight datasets from six open-access studies with varying degrees of movement intensities using increasingly conservative sample rejection criteria. We evaluated the subsequent decomposition quality in terms of the component mutual information, the amount of brain, muscle, and “other” components, the residual variance of the brain components, and an exemplary signal-to-noise ratio.

**Main results:** We found that increasing movements of participants led to decreasing decomposition quality for individual datasets but not as a general trend across all movement intensities. The cleaning strength had less impact on decomposition results than anticipated, and moderate cleaning of the data resulted in the best decompositions.

**Significance:** Our results indicate that the AMICA algorithm is very robust even with limited data cleaning. Moderate amounts of cleaning such as 5 to 10 iterations of the AMICA sample rejection with 3 standard deviations as the threshold will likely improve the decomposition of most datasets, irrespective of the movement intensity.

## 1 Introduction

Removing artifacts from electrophysiological data in the time-domain can be a task as time-consuming as it is important. Rejecting periods of “bad” data that should not be taken into account for further downstream analysis has become a staple in electroencephalography (EEG) analysis from the outset of the method. This includes the rejection of bad epochs when computing event-related measures, but also the rejection of bad samples before running an independent component analysis (ICA; Bell & Sejnowski, 1995; Hyvärinen et al., 2001). ICA is a linear model to decompose the acquired sensor data into components that can subsequently be interpreted regarding the underlying physiological processes (e.g. brain, eyes, muscle, other), and as a common preprocessing step before running ICA, bad samples are removed from the data. Similar to applying a high-pass filter before ICA and copying the decomposition results back to unfiltered data (Klug & Gramann, 2021; Winkler et al., 2015), the ICA results computed on a dataset that had bad time points removed can be applied to the complete uncleaned data in the end (e.g. Gramann et al., 2021; Jacobsen et al., 2021) to retain as much data as possible for downstream analyses. Although time-domain cleaning is regularly used, to our knowledge no study has yet investigated the effect of time-domain cleaning on ICA decomposition in depth. The present study addresses this issue and systematically investigates the effect of time-domain cleaning on the resultant ICA decomposition of real-world data while taking different experimental protocols into account that increase in mobility from stationary to Mobile Brain/Body Imaging (MoBI; Gramann et al., 2011, 2014; Makeig et al., 2009) setups. The goal of this study is to provide best practices that can inform decisions regarding data cleaning of stationary and mobile experiments before applying ICA.

### 1.1 Cleaning of data from mobile experiments is a complex problem

While data cleaning in the time-domain is important for stationary, seated, experiments, it is even more relevant for experiments collecting data from mobile participants. These mobile EEG (Debener et al., 2012) and MoBI (Jungnickel et al., 2019) studies are gaining popularity. Especially with analytical options to remove non-brain activity from high-density EEG data, this approach allows for imaging the human brain in its natural habitat – in participants moving in and actively engaging with their environment. This increased mobility naturally comes with increased non-brain data, traditionally considered artifacts, contributing to the recorded signal.

This non-brain data can be categorized in three groups: 1) Increased physiological activity stemming from the eyes and muscles will be present in the recordings (Gramann et al., 2011; Jungnickel et al., 2019). 2) Movement of, or pressure on the electrodes can lead to mechanical artifacts such as large, transient spikes, and cable sway can result in slow, but high amplitude oscillatory artifacts (Gramann et al., 2011; Oliveira et al., 2017). 3) Electrical artifacts e.g. from line noise or other equipment such as a virtual reality head-mounted display (VR HMD) can be present in the data, and the electrical fields can vary in orientation (respective to the head) and amplitude due to the movement.

In traditional seated experimental protocols, all these contributions to the recorded signal would be candidates for removal. However, as eye and muscle activity can be found throughout the recordings in stationary as well as mobile EEG data, and they typically can be removed with ICA, it is not always clear or easy to decide which time points to remove during time-domain cleaning. Considering mechanical artifacts, large spikes from electrode shifts can be detected comparably easily but cable sway, for example, although potentially high in amplitude, is not a clear case for removal since it might be present in the entire experiment and/or especially in times that are interesting in the experimental paradigm. Taken together, mobile EEG protocols complicate the time-domain cleaning of electrophysiological experiments and traditional heuristics for data cleaning can not always be applied.

### 1.2 Different automatic cleaning options - different challenges

Manually removing samples from data is a sub-optimal approach, as it is both time-consuming and subjective. With the varying experience of the persons cleaning the data, the resulting cleaning strategy will also vary. Even within the same person, different mental states or varying noise levels in different datasets may alter the cleaning procedure. To ensure a reliable, repeatable, and transparent cleaning, it is thus preferred to make use of automatic cleaning algorithms. Several such options exist, ranging from methods based on simple amplitude criteria to the identification of artifactual time periods based on the spectral characteristics to more complex approaches that identify artifactual time periods based on artifact subspaces such as the EEGLAB (Delorme & Makeig, 2004) *clean_rawdata* function, which uses Artifact Subspace Reconstruction (ASR; Kothe & Jung, 2015).

These methods do have their challenges, though. Identifying bad time points by their amplitude alone will either be a very lax measure or it will also remove all periods containing eye blinks since these are high-amplitude signals. Removing eye-blinks before ICA decomposition, however, is not desired, as ICA can typically remove these more reliably and preserve the respective time points for downstream analyses. Spectral measures will be prone to removing periods with muscle activity since these are usually detected by having increased high-frequency broad-band power (Onton & Makeig, 2006). But especially in mobile experiments, removing time periods with muscle activity would result in excessive cleaning, and the computed ICA decomposition would not be readily applicable to the entire dataset since it was not informed by time points containing muscle activity. And while the cleaning threshold of ASR can be adjusted to remove mainly large transient spikes, ASR is very sensitive to this threshold (Chang et al., 2019), and especially for mobile data, it does not always find a suitable baseline by itself. ASR thus requires a specifically recorded baseline and sometimes different cleaning thresholds for different movement modalities and even different datasets within the same modality, which renders it unsuitable for automatic data cleaning as targeted in this study.

### 1.3 AMICA sample rejection

The Adaptive Mixture ICA (AMICA; Palmer et al., 2011), currently one of the most powerful ICA algorithms (Delorme et al., 2012), includes an inbuilt function to reject bad samples that might not be well-utilized by researchers working with the algorithm: AMICA can reject bad samples based on their log-likelihood while computing the decomposition model. The log-likelihood is an objective criterion corresponding to the algorithm’s estimate of the model fit, effectively leading to the rejection of samples AMICA cannot easily account for. Hence, unlike other cleaning methods, this option will only remove those kinds of artifacts that negatively affect the decomposition and retain those that can be decomposed and removed with ICA. This is done in an iterative fashion: First, several steps of the model estimation are performed, then samples are rejected based on the difference of their log-likelihood from the mean in standard deviations (SDs), then the model is estimated for several steps again before the next rejection, and so on. The start of the rejection, the number of rejection iterations, the SD threshold as well as the number of model computation steps between each rejection can be set in the AMICA computation parameters. This artifact rejection approach is model-driven and allows users to automatically remove time-domain data to improve the decomposition.

When applied from the EEGLAB user interface (AMICA plugin v1.6.1), this is disabled by default, but when opening the rejection sub-interface, it is enabled with 5 iterations with 3 SDs, starting after the first AMICA step, with one step between each iteration. When the *runamica15* function is applied directly from the command line, the cleaning is disabled by default, but when enabled, it rejects three times with 3SDs, starting after the second AMICA step, with 3 steps between each iteration. As a consequence, different settings will impact whether and how AMICA uses time-domain cleaning during the decomposition. However, while previous evaluations have shown that AMICA is one of the currently best algorithms for EEG decomposition (Delorme et al., 2012), the impact of the integrated time-domain cleaning procedure has not been evaluated yet.

### 1.4 Current study

In this study, we thus investigate the impact of automatic sample rejection on the quality of the AMICA decomposition of data from six experiments with different levels of mobility. To this end, we varied the cleaning intensity in terms of the number of cleaning iterations as well as the rejection threshold. As measures of decomposition quality, we used the number of brain, muscle, and unspecified components, the residual variance of brain components, and the component mutual information. Additionally, we examined the signal-to-noise ratio in one standing and one mobile condition of the same experiment. We hypothesized that 1) increasing mobility affects the decomposition negatively, 2) cleaning affects the decomposition positively, and 3) an interaction exists, where experiments with more movement require more cleaning to improve their decomposition quality. We had no hypothesis as to what the optimal amount of cleaning should be. Based on the empirical evidence from this study, we formulate recommendations for using sample rejection with AMICA.

## 2 Methods

### 2.1 Datasets

For a reliable estimate of the effect of time-domain cleaning on the quality of the ICA decomposition for datasets from stationary as well as mobile EEG protocols, we included open access EEG datasets with a wide range of movement conditions in this study. We used datasets that used at least 58 EEG channels (not including EOG), had a sampling rate of at least 250 Hz, and either contained channel locations or standard extended 10-20% system electrode layouts. Representation of different EEG setups was ensured by including no more than two datasets from one lab. We manually subsampled channels where necessary as described in the section *Preparation*. This resulted in eight datasets from six studies containing standard seated protocols but also a gait protocol, arm reaching, and irregular movement protocols.

The effect of movement on the EEG is not straightforward as it does not only depend on the general amplitude of movement, but more so on the acceleration and the proximity to the electrodes. We thus categorized the available datasets into three groups: 1) Seated or standing tasks with limited head movement (low intensity): While some small artifacts might still be present, these types of experiments are expected to exhibit few artifacts other than eye movement and minimal neck muscle activity for stabilization. 2) Seated or standing tasks with head movement (medium intensity): These types of experiments are expected to exhibit increased physiological artifacts from the neck and eyes, as well as some mechanical artifacts due to small electrode shifts during the movement. 3) Walking and other high-amplitude physical movements (high intensity): These types of experiments are expected to exhibit a large amount of head movement, including eye movement as well as neck muscle activity for head stabilization and balance. Gait can also lead to cable sway and oscillations in pressure of the electrodes, leading to additional, potentially strong, mechanical artifacts.

#### Video Game

This dataset is available at https://doi.org/10.18112/openneuro.ds003517.v1.1.0 (Cavanagh & Castellanos, 2021) and contains data of 17 participants (6 female and 11 male, mean age = 20.94 years, SD = 5.02 years). Participants were sitting while playing a video game using a gamepad. Data was recorded with a 500 Hz sampling rate using 64 electrodes (Brain Products GmbH, Gilching, Germany), and filtered with a high-pass filter of .01 Hz and a low-pass filter of 100 Hz. Channels were manually downsampled by selecting 58 channels of only scalp electrodes. This dataset was categorized as having low movement intensity due to the limited amount of head movement.

#### Face Processing

This dataset is available at https://doi.org/10.18112/openneuro.ds002718.v1.0.5 (Wakeman & Henson, 2021) and contains data of 19 participants (8 female and 11 male, age range 23- 37 years). Participants were seated in front of a screen and exposed to images of faces. Data was recorded with a 1100 Hz sampling rate using 70 electrodes and a 350 Hz low-pass filter was applied. Channels were manually downsampled by selecting 65 channels that were closest to corresponding electrodes from an extended 10-20% layout of scalp electrodes. This dataset was categorized as having low movement intensity due to the limited amount of head movement.

#### Spot Rotation (stationary/mobile)

This dataset is available at https://doi.org/10.14279/depositonce-10493 (Gramann et al., 2021) and contains data of 19 participants (10 female and 9 male, aged 20–46 years, mean age = 30.3 years). The experiment consisted of a rotation on the spot, which either happened in a virtual reality environment with physical rotation or in the same environment on a two-dimensional monitor using a joystick to rotate the view. Participants were standing in front of the computer screen in the stationary condition. The data was split into the two conditions of joystick rotation (stationary) and physical rotation (mobile) for the purpose of this study. Data was recorded with a 1000 Hz sampling rate using 157 electrodes (129 on the scalp in a custom equidistant layout, 28 around the neck in a custom neckband) with the BrainAmp Move System (Brain Products GmbH, Gilching, Germany). Channels were manually downsampled by selecting 60 channels that were closest to corresponding electrodes from an extended 10-20% layout of scalp electrodes. This dataset was recorded in the same laboratory as the *Prediction Error* dataset. The stationary dataset was categorized as having low movement intensity due to the limited amount of head movement.

The mobile dataset involved a physical, but slow rotation on the spot, including head movement, and was thus categorized as having medium movement intensity.

#### Prediction Error

This dataset is available at https://doi.org/10.18112/openneuro.ds003846.v1.0.1 (Gehrke et al., 2021) and contains data of 20 participants (12 female, mean age = 26.7 years, SD = 3.6 years) of which one was removed by the authors due to data recording error. Participants were seated at a table and equipped with an HMD. The task consisted in reaching for virtual cubes that appeared in front of participants on the table. Participants moved their arm and upper torso to reach the virtual goal on the table. Data was recorded with a 1000 Hz sampling rate using 64 electrodes (Brain Products GmbH, Gilching, Germany). Channels were manually downsampled by selecting 58 channels of only scalp electrodes. This dataset was recorded in the same laboratory as the *Spot Rotation* dataset. This dataset was categorized as having medium movement intensity due to the involved head movement during the reaching task.

#### Beamwalking (stationary/mobile)

This dataset is available at https://doi.org/10.18112/openneuro.ds003739.v1.0.2 (Peterson & Ferris, 2021) and contains data of 29 participants (15 female and 14 male, mean age = 22.5 years, SD = 4.8 years). Participants either stood or walked on a balance beam and were exposed to sensorimotor perturbations. Perturbations were either virtual-reality-induced visual field rotations or side-to-side waist pulls. Because of the different degrees of movement, the data was split according to the two conditions: stationary (standing) and mobile (walking). Data was recorded at a 512 Hz sampling rate using 136 electrodes (BioSemi Active II, BioSemi, Amsterdam, The Netherlands). Channels were manually downsampled by selecting 61 channels that were closest to corresponding electrodes from an extended 10-20% layout of scalp electrodes. The stationary dataset involved head movement due to the disturbances of the balance and was thus categorized as having medium movement intensity. The mobile dataset additionally involved slow walking, and higher-acceleration movements can be expected from balance perturbations during walking. It was thus categorized as having high movement intensity

#### Auditory Gait

This dataset is available at https://doi.org/10.1038/s41597-019-0223-2 (Wagner et al., 2019) and contains data of 20 participants (9 females and 11 males, aged 22–35 years, mean age = 29.1 years, SD = 2.7 years). Participants had to walk on a treadmill and synchronize their steps to a regular auditory pacing stream that included infrequent, sudden shifts in tempo. Data was recorded with a 512 Hz sampling rate using 108 electrodes with seven 16-channel amplifiers (g.tec GmbH, Graz, Austria), high pass filtered >0.1 Hz, low pass filtered <256 Hz and a notch filter was applied at 50 Hz to remove power line noise. Channels were manually downsampled by selecting 61 channels that were closest to corresponding electrodes from an extended 10-20% layout of scalp electrodes. This dataset was categorized as having high movement intensity since it involved gait on a treadmill.

### 2.2 Data processing

All data was processed in an automated fashion with identical preprocessing steps as displayed in figure 1. The main processing steps can be summarized under pre-processing, AMICA with sample rejection, and ICA post-processing, followed by the computation of quality measures to evaluate the decomposition.

**Figure 1:**
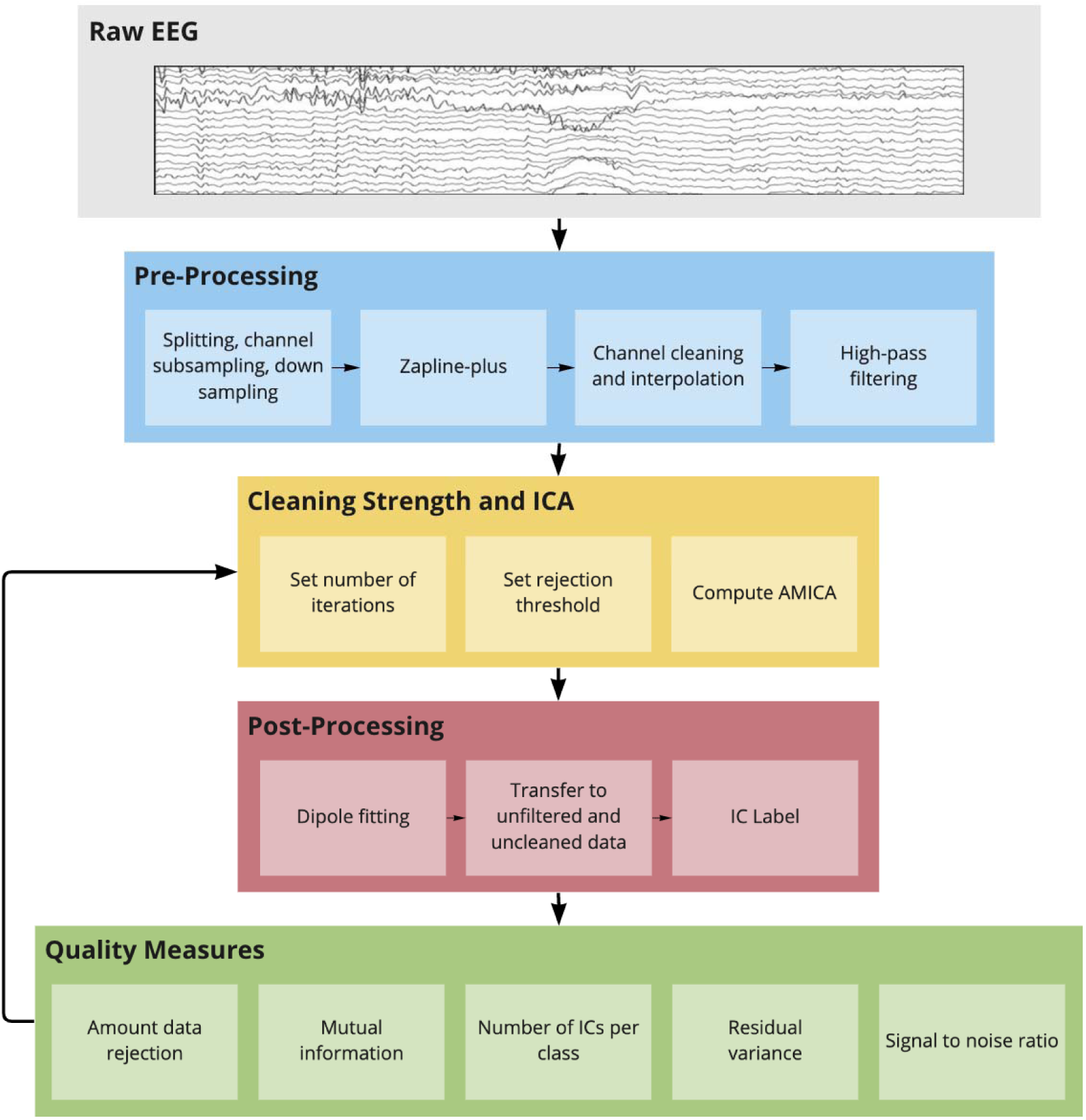
Data processing pipeline. The number of cleaning iterations and the rejection threshold were varied several times to compute AMICA and post-processing as well as quality measures repeatedly. Details can be found in section *Data processing*.

#### 2.2.1 Preparation

All datasets were first loaded into EEGLAB, and if a study contained data of two different conditions these were split and subsequently treated separately. We then manually selected channels to reduce the number of channels to a range of 58 to 65, excluding EOG or neck channel locations, and matched the remaining channels to the extended 10-20% layout as close as possible if the original layout was equidistant (see section *Datasets*). We did not subsample all studies to exactly 58 channels to allow an evenly distributed whole head coverage in all datasets. Full comparability of the channel layout between studies was not given, nor was it intended since the data were recorded in different labs with different devices. In addition, a previous investigation revealed a ceiling effect in obtaining brain ICs with an increasing number of electrodes used for ICA decomposition (Klug & Gramann, 2021). A doubling of channels from 64 to 128 resulted in only 3 to 4 more brain ICs while differences in the number of brain ICs due to different movement intensities were more substantial with around 5 to 6 brain ICs. As a consequence, we expected our differences in channel count to have minimal impact on the computed quality measures. After channel reduction, all datasets were downsampled to 250 Hz and reduced to a length of 10 minutes (150,000 samples), ensuring there was the same amount of data available for all datasets. All data were subsequently processed using the *BeMoBIL pipeline* (Klug et al., 2022) with identical parameters except for varying time-domain cleaning types and strengths.

#### 2.2.2 Zapline-plus

The *Auditory Gait* dataset had a notch filter applied before uploading. All other datasets were processed with Zapline-plus (de Cheveigné, 2020; Klug & Kloosterman, 2022) to remove line noise and other frequency-specific noise peaks. Zapline (de Cheveigné, 2020) removes noise by splitting the data into an originally clean (data A) and a noisy part (data B) by filtering the data once with a notch filter (A) and once with the inverse of this notch filter (B). It then uses a spatial filter to remove noise components in the noisy part (B) to get a cleaned version of that part (B’). Finally, the two clean parts (A and B’) are added back together to result in a cleaned dataset with full rank and full spectrum except for the noise. Zapline-plus is a wrapper for Zapline that chunks the data to improve the cleaning and adapts the cleaning intensity automatically, thus maximizing the cleaning performance while ensuring minimal negative impact. We used default parameters in all datasets containing mobile data (*Spot Rotation, Prediction Error, Beamwalking*), but limited the noise frequency detector to line noise for the *Face Processing* and *Video Game* datasets to avoid the removal of minor spectral peaks in stationary datasets that were not expected to be influenced by additional electronic equipment or mechanical artifacts.

#### 2.2.3 Channel cleaning and interpolation

We detected bad channels using the *clean_rawdata* plugin of EEGLAB in an iterative way. We did not use the *flatline_crit*, the *line_noise_crit*, and samples were not rejected, nor was ASR applied. The only used criteria to detect bad channels was the *chancorr_crit* criterion, which interpolates a channel based on a random sample consensus (RANSAC) of all channels and then computes the correlation of the channel with its own interpolation in windows of 5 s duration. If this correlation is below the threshold more than a specified fraction of the time (50% in our case), it is determined to be bad. Since the function has a random component, it does not necessarily result in a stable rejection choice, which is why this detection was repeated ten times, and only channels that were flagged as bad more than 50% of the time were finally rejected. Removed channels were then interpolated and the data was subsequently re-referenced to the average using the *full rank average reference* plugin of EEGLAB, which preserves the data rank while re-referencing.

#### 2.2.4 High-pass filtering

High-pass filtering has a positive effect on the ICA decomposition quality and is especially important in mobile studies (Klug & Gramann, 2021). For a dataset with 64 channels, a filter of 0.5 to 1.5 Hz cutoff resulted in the best decomposition for both stationary and mobile conditions (Klug & Gramann, 2021), which is why we chose a cutoff of 1 Hz in this study. We specified the filter manually as recommended (Widmann et al., 2015) and used the same filter specifications as in Klug & Gramann (2021): a zero-phase Hamming window FIR-filter (EEGLAB firfilt plugin, v1.6.2) with an order of 1650 and a passband-edge of 1.25 Hz, resulting in a transition bandwidth of 0.5 Hz and a cutoff frequency of 1 Hz.

#### 2.2.5 Independent component analysis with sample rejection

All final datasets were decomposed using AMICA with different numbers of sample rejection iterations and different rejection thresholds to compare the results of the decomposition for the eight datasets. We used one model and ran AMICA for 2000 iterations. Since we interpolated channels previously we also let the algorithm perform a principal component analysis rank reduction to the number of channels minus the number of interpolated channels. As we used the full rank average reference, we did not subtract an additional rank for this. All computations were performed using four threads on machines with identical hardware, an AMD Ryzen 1700 CPU with 32GB of DDR4 RAM.

In this step, we investigated the effect of different cleaning intensities using the AMICA sample rejection algorithm. All rejection was started after the *runamica15* default 2 iterations, with the default 3 iterations between rejections. We repeated the AMICA computation process either without sample rejection, or with 1, 3, 5, 7, or 10 iterations using 3 SDs as the threshold, and additionally with 10, 15, and 20 iterations using 2.8 SDs, and last with 20 iterations using 2.6 SDs.

#### 2.2.6 Dipole fitting

An equivalent dipole model was computed for each resulting independent component (IC) using the DIPFIT plugin for EEGLAB with the 3-layer boundary element model of the MNI brain (Montreal Neurological Institute, Montreal, QC, Canada). The dipole model includes an estimate of IC topography variance which is not explained by the model (residual variance, RV). For datasets that had individually measured electrode locations of an equidistant channel layout, the locations were warped (rotated and rescaled) to fit the head model.

#### 2.2.7 Transfer of ICA to unfiltered data

One of the goals of this study was to investigate the effect of time-domain cleaning on the component mutual information (MI) and the signal-to-noise ratio (SNR) when applied to the full dataset. To this end, the resulting AMICA decomposition and dipole models were copied back to the preprocessed dataset from section *Channel cleaning and interpolation* (line noise removed, channels interpolated, re-referenced to the average, but no high-pass filter and no time-domain cleaning).

#### 2.2.8 Independent component classification using ICLabel

In order to categorize the ICs according to their likely functional origin, we applied the ICLabel algorithm (Pion-Tonachini et al., 2019). ICLabel is a classifier trained on a large database of expert labelings of ICs that classifies ICs into brain, eye, muscle, and heart sources as well as channel and line noise artifacts and a category of other, unclear, sources. As it was shown that the ‘lite’ classifier worked better than the ‘default’ one for muscle ICs (Klug & Gramann, 2021), we used the ‘lite’ classifier in this study. We used the majority vote to determine the final class, meaning the IC received the label with the highest probability.

#### 2.2.9 Quality Measures

To measure the impact of the cleaning on the ICA decomposition, we used several measures that addressed both mathematical and practical considerations of the ICA decomposition: i) The mutual information (MI) of the components after applying the ICA solution to the complete dataset. The MI is essentially the mathematical description of how well the ICA can decompose the data, as the ICA minimizes component MI. ii) The number of ICs categorized as stemming from *brain*, *muscle*, and *other* sources defined by ICLabel, as especially in MoBI research not only brain, but all physiological sources can be of interest to the experimental analysis. iii) The mean RV of brain ICs, as this can be considered a measure of the physiological plausibility of the IC (Delorme et al., 2012). iv) The combination of the above measures as the ratio of the number of brain ICs by the mean brain RV (higher indicates better decomposition), v) An exemplary computation of the signal-to-noise ratio (SNR) on the two *Spot Rotation* datasets (physical rotation in VR and 2D monitor rotation), using the same measures as in (Klug & Gramann, 2021). For this, we removed all non-brain ICs in the final dataset and computed event-related potentials (ERPs) of the trial onsets at the electrode closest to the Pz electrode in the 10-20% system. On average, 30.58 (SD = 7.31) epochs were used per subject and condition. This comparably low number of epochs is caused by the previous reduction in dataset length to 10 minutes, and the results of this approach should be interpreted with care. The signal was defined as the mean amplitude from 250 ms to 450 ms and the standard deviation in the 500 ms pre-stimulus interval was used as a measure of the noise (Debener et al., 2012).

### 2.3 Statistical Analysis

To quantitatively assess the influence of time cleaning on data quality, we used a linear mixed effects model (LMM) with the following design:

data_quality_measure ∼ movement_intensity * cleaning_intensity + (1 | study/subject)

The fixed effects included movement intensity and cleaning intensity, as well as their interaction. Movement intensity was modeled as an ordered factor with the categories “high”, “medium”, and “low”, with the classifications outlined in Section 2.1. Successive difference contrasts were used for post-hoc analysis to assess differences between these levels. Cleaning intensity, on the other hand, was modeled as a continuous variable ranging from 0 to 9, corresponding to the lowest level (no cleaning) to the highest level (20 iterations, sigma = 2.6). Although cleaning intensity is not technically continuous, we chose to model it in this way because a discrete variable with ten levels would sacrifice a significant amount of statistical power, resulting in an inflation of type-II statistical error. We considered studies and subjects as random factors. The random factor “subject” was nested within “study” because we expected variation in the study due to factors such as the equipment used and data collection practices, as well as an effect of subjects beyond that.

In an additional attempt to investigate the legitimacy of our classification of movement intensity, we opted for a data-driven clustering of studies based on the AMICA quality measures when no cleaning was applied. A Gaussian Mixture Model (GMM) was used as the clustering algorithm. We set the number of mixture components to three to allow comparability with the a priori study classification. The same LMMs were then recalculated using the movement intensity categorization of the new clustering solution, and the resulting models were compared with the previous ones to determine whether they provided a better explanation of the data. To this end, we used the Akaike information criterion (AIC), which indicates the relative quality of statistical models. A difference in AIC greater than 10 is considered to be strong support that the model with the lower AIC fits the data better (Burnham & Anderson, 2004). Since the data-driven clusters lacked inherent order of low-mid-high intensity and we were only interested in the general model fit, no post hoc analysis was performed.

In addition, another set of LMMs was estimated for the *Spot Rotation* and *Beamwalking* studies in a within-study design of the movement intensities. Here, the movement intensity factor had only ’low’ and ’high’ values, corresponding to the stationary and mobile conditions, respectively. Successive differences were again used for the post hoc analysis.

For the exemplary SNR analysis, we computed the previous model used for the *Spot Rotation*, excluding the “no ICA” level. To examine the impact of the ICA on the SNR, we computed another LMM incorporating only the “no ICA” level and the “ICA no time-cleaning” level using the following formula:

snr ∼ movement_intensity * ICA + (1 | subject)

## 3 Results

We anticipated three effects regarding the time-domain cleaning and movement intensity on the decomposition quality of the ICA. First, we expected a higher movement intensity to decrease the decomposition quality. Second, higher cleaning intensity should remove more artifacts and therefore result in a better decomposition quality. Finally, we expected an interaction of movement intensity and cleaning intensity such that more movement would require more cleaning to reach a better decomposition.

### 3.1 The effect of movement intensity on the decomposition quality

The most fundamental effect of changing the hyperparameters of AMICA sample rejection is the amount of data that is rejected. If more movement results in more problematic artifacts (those that AMICA can not easily include in its model e.g. due to nonlinearities), more movement should also result in more rejected samples. As can be seen in figure 2a, the dataset with the lowest amount of rejection was indeed a set with low movement intensity, namely the *Face Processing* dataset. Yet, the dataset with the second lowest removal was a high movement intensity set (*Auditory Gait*). Furthermore, the most data was rejected in the *Spot Rotation (stationary)* set, which was a set with low movement intensity. In our results, movement intensity did not show a significant main effect, neither between the low and medium groups (β=0.09, SE=1.29, t=0.07, p=0.94) nor between the medium and high groups (β=-0.49, SE=1.42, t=-0.35, p=0.74).

**Figure 2:**
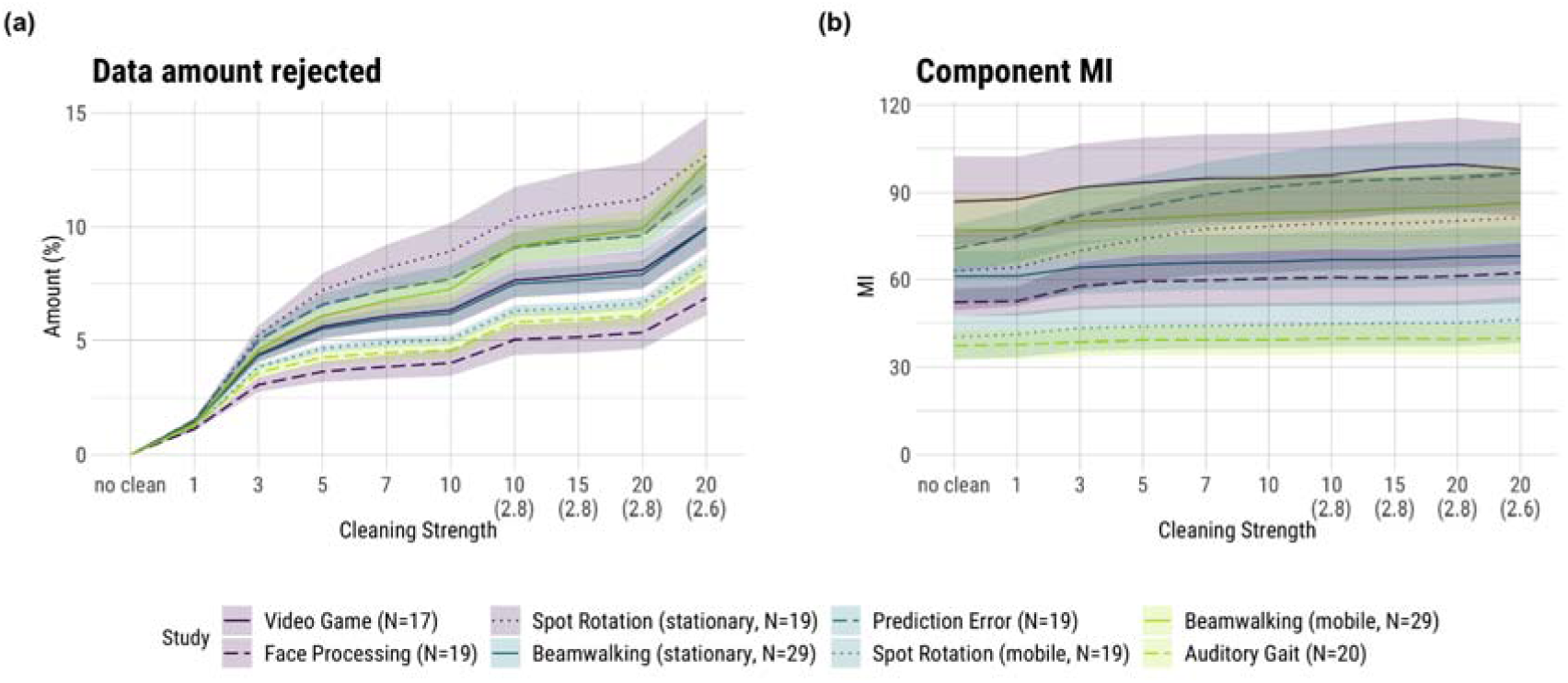
Results for the data rejection amount (a) and component mutual information (MI; b). Shaded areas depict the standard error of the mean (SE). The numbers on the abscissa refer to the number of iterations of the sample rejection, with a default of 3 SDs as the rejection threshold. The numbers in brackets on the abscissa refer to the rejection threshold in SDs when deviating from the default. “No clean” refers to no sample rejection being applied when computing AMICA. The colors denote the movement intensities: purple - low, blue - medium, green - high.

Figure 2b shows the results for the MI, where the AMICA reached the lowest MI, and thereby its best decomposition from a mathematical point of view, in the *Auditory Gait* and *Spot Rotation (mobile)* datasets. The *Auditory Gait* and *Spot Rotation (mobile)* datasets were categorized as high and medium-to-high movement intensity, respectively. Even when comparing movement conditions of the same studies, no clear trend was identifiable. While the results for the *Beamwalking* study yielded a higher MI in its mobile condition, the *Spot Rotation* study showed the opposite trend.

Statistical analysis revealed no significance in both the main effect of the low and medium groups (β=-9.35, SE=18.34, t=-0.51, p=0.63) and the medium and high groups (β=-1.38, SE=20.08, t=-0.07, p=0.95). Of note, the intraclass correlation coefficient (ICC) of this model is 0.98, indicating that the variance in MI is almost entirely accounted for by the random effects of between-subject and between-study variability.

The results of the number of resulting brain and muscle ICs can be seen in figure 3a. The highest amounts of brain ICs were found in the datasets *Face Processing*, *Beamwalking (stationary)*, and *Auditory Gait*, with the last dataset having high movement intensity. The group of datasets with the lowest number of brain ICs (with around 6 fewer than the highest) consisted of two medium movement intensity studies but also included the low movement intensity *Video Game* study. This pattern of ambiguity is also reflected in the statistical analysis. The main effect for the low to medium group (β=-0.79, SE=2.70, t=-0.29, p=0.78) contrasts with the effect for the medium to high group (β=2.53, SE=2.99, t=0.85, p=0.44). Both main effects are not significant. It is also important to note the high intraclass correlation coefficient (ICC) of 0.92.

**Figure 3:**
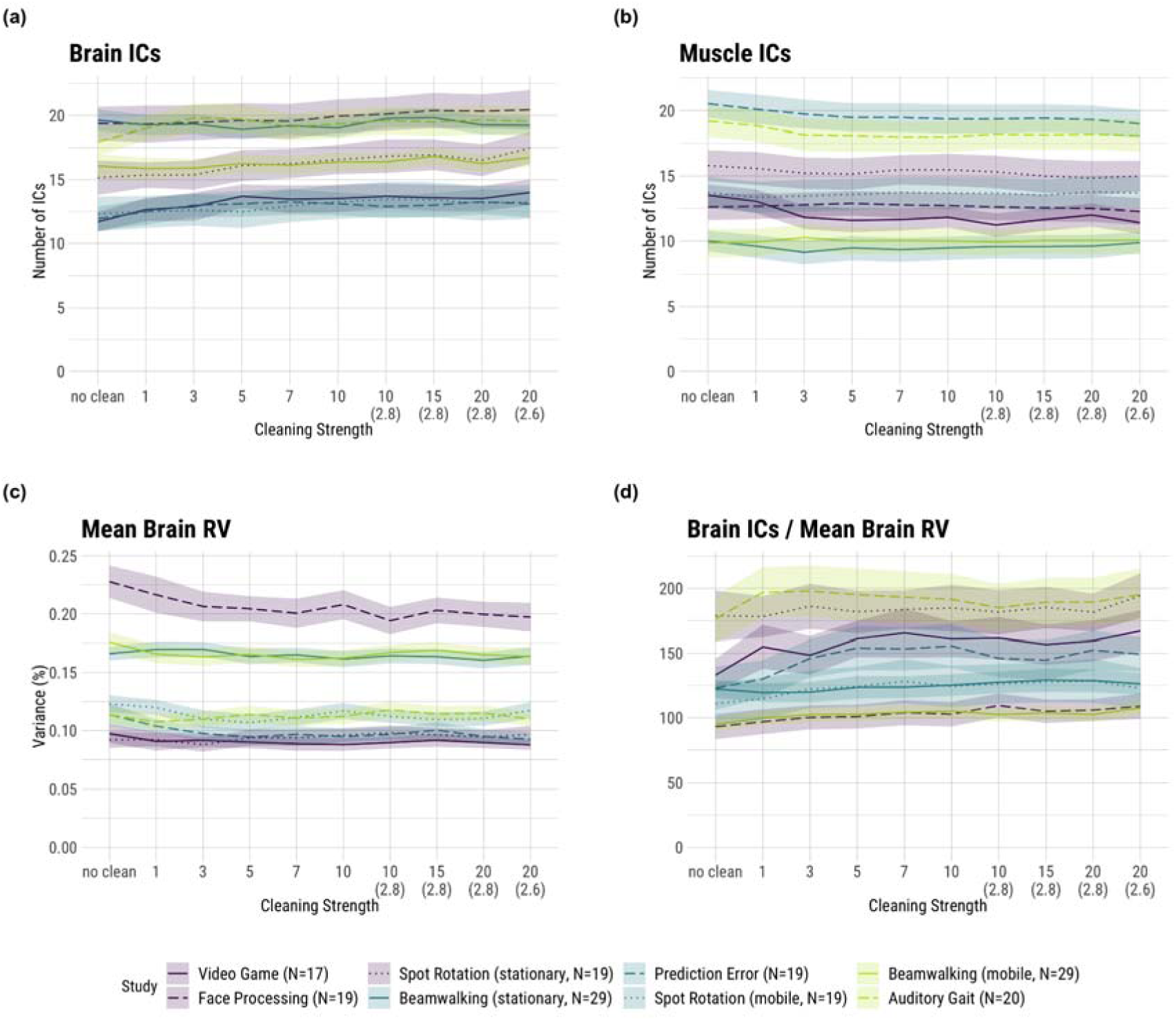
Results for the brain (a) and muscle (b) ICLabel classifications, residual variance (RV; c) and the ratio of the number of brain ICs to their mean RV (d). Shaded areas depict the standard error of the mean (SE). The numbers on the abscissa refer to the number of iterations of the sample rejection, with a default of 3 SDs as the rejection threshold. The numbers in brackets on the abscissa refer to the rejection threshold in SDs when deviating from the default. “No clean” refers to no sample rejection being applied when computing AMICA. The colors denote the movement intensities: purple - low, blue - medium, green - high.

The number of muscle ICs also did not show a clear trend with movement intensity: one of the datasets with the least amount of muscle ICs was the high movement intensity set *Beamwalking (mobile)*. This qualitative description is further supported by the statistical analysis, which showed main effects of contrasting signs (low to medium: β=0.61, SE=3.52, t=0.17, p=0.87; medium to high: β=-0.10, SE=3.91, t=-0.02, p=0.98). Both effects were found to be nonsignificant. The ICC for this model was 0.98.

As can be seen in figure 3c, the highest RV values (indicating lowest physiological plausibility) for brain ICs (above 20%) were found in the *Face Processing* set, which was a low movement intensity study. The three datasets of *Prediction Error*, *Spot Rotation (stationary)*, and *Video Gaming*, coming from two different movement intensity groups, formed the cluster with the lowest RVs of around 10%. Taken together, the movement intensity of a study had no clear effect on the quality of the brain ICs. This is further supported by the fact that the best and worst ratio of the number of brain ICs to their mean RV was attained by two studies of the high movement intensity group. The statistical analysis showed no significant main effects (low to medium, β=-4.03e-03, SE=4.07e-02, t=- 0.10, p=0.92; medium to high, β=1.03e-02, SE=4.55e-02, t=0.23, p=0.83). The ICC of this model was 0.93.

Figure 4a shows the number of ICs labeled as “other”. The highest numbers of “other” ICs were attained by the datasets from the low-intensity group followed by the medium movement intensity group. The *Prediction Error* set, belonging to the medium movement intensity group, had 17-18 “other” ICs and therefore around 5 “other” ICs fewer than the rest of the medium movement intensity group. The lowest number of “other” ICs had the high movement intensity study *Auditory Gait* reaching fewer than 15 “other” ICs. Not aligned with this trend were the *Spot Rotation (mobile)* and the *Beamwalking (mobile)* datasets stemming from the medium and high movement intensity group, respectively, but having similar amounts of “other” ICs as the low cluster. The statistical analysis showed no significant main effects (low to medium, β=-2.68, SE=2.97, t=-0.91, p=0.41; medium to high, β=-2.87, SE=3.29, t=-0.87, p=0.42). The ICC for this model was 0.92.

**Figure 4:**
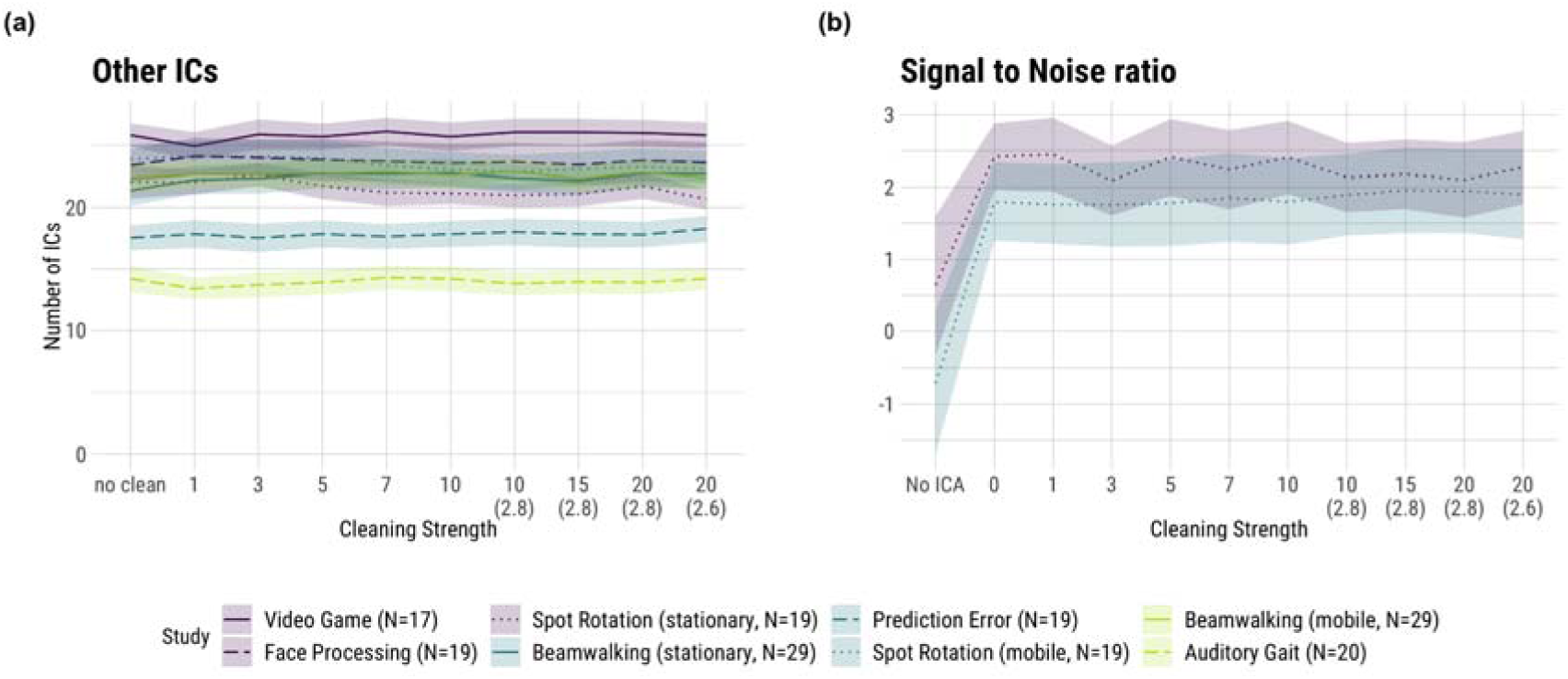
Results for other ICs (a) and exemplary signal-to-noise ratio (SNR; b). The SNR was computed on the *Spot Rotation* study only. Shaded areas depict the standard error of the mean (SE). The numbers on the abscissa refer to the number of iterations of the sample rejection, with a default of 3 SDs as the rejection threshold. The numbers in brackets on the abscissa refer to the rejection threshold in SDs when deviating from the default. “No clean” (left plot) / “0” (right plot) refers to no sample rejection being applied when computing AMICA. “No ICA” (right plot) refers to the SNR values being computed on the raw dataset without any ICA cleaning. The colors denote the movement intensities: purple - low, blue - medium, green - high.

**Figure 5:**
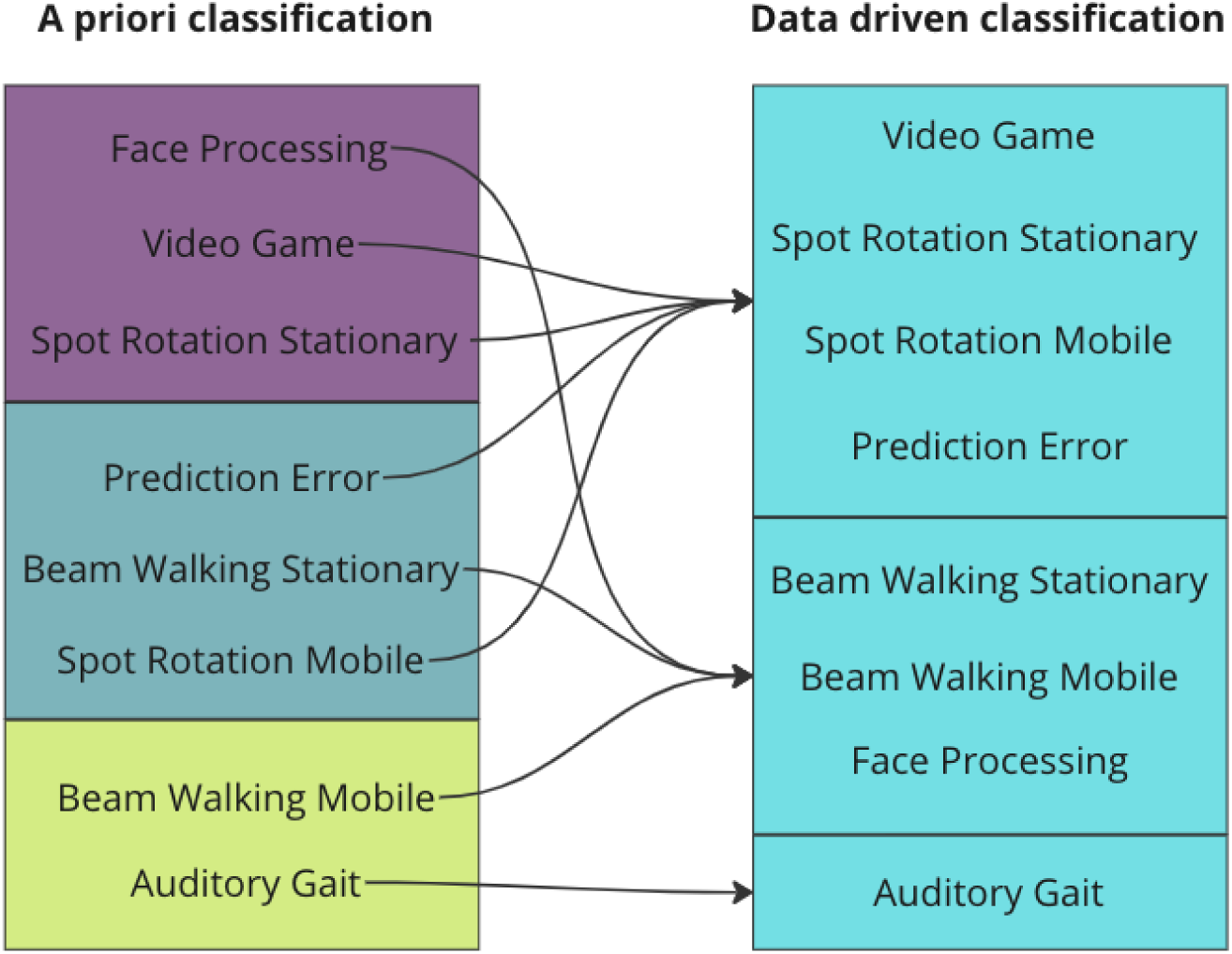
Differences in clustering of studies comparing the a priori classification with the methodology described in section X. The colors on the left hand side denote the movement intensities: purple - low, blue - medium, green - high. The right hand side has no color scheme because the clustering has no inherent ordering.

Lastly, in many cases, ICA is used as a means for data cleaning or extracting specific aspects of the signal. Hence, we used the SNR of an ERP in one exemplary study, *Spot Rotation*, to give a practical example of the effect of the use of ICA. As can be seen in figure 4b, this example revealed a generally lower SNR in the mobile condition. However, this effect of mobility was not significant, as confirmed by both the 2x2 model that did not include cleaning strength, but the use of ICA in general (β=-0.99, SE=0.69, t=-1.42, p=0.16) and the model that tested the effect of the cleaning strength, although the latter almost showed an indication of an effect (β=-0.67, SE=0.34, t=-1.96, p=0.051).

### 3.2 The effect of cleaning intensity on the decomposition quality

Increasing the number of cleaning iterations and decreasing the rejection threshold resulted in significantly more samples being rejected (figure 2a; β=1.01, SE=0.02, t=61.77, p<0.01). The effect of cleaning iterations reached a ceiling after around 5 iterations and substantial increases in data rejection were only reached when in addition to more iterations the rejection threshold was lowered as well. As the MI is computed on the entire dataset but the ICA did not take the rejected samples into account, all studies showed a monotonous increase in MI with stronger cleaning (figure 2b). Nevertheless, the magnitude of this effect was only moderate as in most datasets the MI remained almost constant and within the range of their SEs. Two sets (*Prediction Error*, *Spot Rotation (stationary)*) showed a stronger increase in the low iterations but also leveled off after around 7 iterations.

The number of brain ICs did not vary with increased cleaning intensity outside of the SE range for almost all studies (figure 3a). The main effect of cleaning intensity was nevertheless significant with a small effect size (β=0.12, SE=0.01, t=8.66, p<0.01). Only the *Audiocue* set showed a more noticeable increase in low amounts of cleaning (3-5 iterations) of around 2 ICs but this effect reached a ceiling for stronger cleaning. Other sets exhibited small trends in the same direction, but no strong effect could be seen. All datasets except for *Beamwalking (stationary)* exhibited a slight decrease in mean RV with the first few iterations of the cleaning. This trend was within the range of the SE, however, and reached a floor after around 5 to 7 iterations. Similar to brain ICs the main effect of cleaning intensity on RV was statistically significant but with a marginal effect size (β=-6.11e-04, SE=1.4e-04, t=- 4.353, p<0.01) . Combining this small trend with the small trend in the number of brain ICs, we did find a positive effect of time cleaning on the ratio of the number of brain ICs to their mean RV (β=1.21, SE=0.16, t=7.65, p<0.01) (figure 3d). Here, all studies exhibited an increase in this ratio with increased cleaning, up to a point of around 7 to 10 iterations. Especially the datasets *Video Game* and *Spot Rotation (stationary)* showed around a 20% increase in this ratio.

Considering the SNR, no effect of stronger time-domain cleaning was visible in our study (figure 4b). There was a pronounced effect when generally using the ICA as a means to select the signal of interest by removing all non-brain ICs (β=2.14, SE=0.69, t=3.09, p<0.01), but the time cleaning itself did not significantly affect the SNR (β=-0.02, SE=0.04, t=-0.62, p=0.54).

### 3.3 The interaction effect of cleaning intensity and movement intensity on the decomposition quality

The changes in the amount of rejected data when cleaning was intensified were very visually similar for all studies (figure 2a) but we nevertheless found several significant interactions. Studies in the high movement intensity group had a slightly greater increase in the amount of rejected data than those in the medium group (β=0.08, SE=0.04, t=2.03, p=0.04). Looking at the MI, we found that the effect of cleaning intensity was less strong for the high movement intensity compared to the medium movement intensity group (β=-0.57, SE=0.16, t=-3.67, p<0.01). The number of brain ICs also showed a significant interaction. The medium movement intensity group had a significantly weaker increase in the number of brain ICs with increased cleaning compared to the low group (β=-0.12, SE=0.03, t=-3.95, p<0.01). The opposite interaction was observed for the number of muscle ICs. Here, the medium movement intensity group had a smaller decrease in the number of muscle ICs with more cleaning than the low group (β=0.06, SE=0.02, t=3.41, p<0.01). While the low and medium movement intensity groups showed a decrease in mean brain RV with more cleaning, this was not the case for the high movement intensity group (β=7.19e-04, SE=3.40e-04, t=2.12, p=0.03). The same interaction was found for the ratio of brain ICs to RV, where the high group showed a significantly weaker increase than the medium movement intensity group (β=-0.82, SE=0.38, t=-2.12, p=0.03). There were no other statistically significant interactions.

### 3.4 Data driven clustering compared to a priori clustering of studies

The data driven clustering created imbalanced clusters with one cluster only containing the Auditory Gait study, one cluster containing both Beam Walking and the Face Processing studies and one for the other four studies. The clustering did not seem to follow any of the aspects used in the a priori clustering.

The LMMs with the data driven clusters had a significantly lower AIC than the models using the a priori clusters for all quality measures except the number of brain ICs (see table 1). Only the number of brain ICs was better explained by the a priori cluster, but the difference was not significant.

**Table 1:** AIC of LMMs for different quality measures based on different clusters of studies. The a priori cluster LMMs used a previously classified, movement intensity as a fixed effect while the data driven cluster LMMs used a classification by clustering based on data quality before time cleaning. An AIC difference greater than 10 is considered to be significant (Burnham & Anderson, 2004).

| Quality measure | AIC a priori clusters | AIC data driven clusters | Difference |
| --- | --- | --- | --- |
| Mutual Information | 12911.4 | 12874.4 | -37.0 |
| Brain ICs | 7234.0 | 7237.2 | 3.2 |
| Muscle ICs | 5780.9 | 5766.7 | -14.2 |
| Mean Brain RV | -8499.6 | -8512.3 | -12.7 |

### 3.5 Within-study comparisons

When looking at the within-study differences, the mobile conditions in both the *Spot Rotation* and the *Beamwalking* study had around 3 brain ICs fewer than their respective stationary counterparts (β=-3.18, SD=0.34, t=-9.41, p<0.01). This was not the case for the number of muscle ICs, however (β=-0.54, SD=0.32, t=-1.70, p=0.09). The brain RV values were significantly higher in the mobile conditions (β=0.01, SD<0.01, t=3.45, p<0.01).

The cleaning intensity had a significant effect only for the number of brain ICs (β=0.10, SD=0.03, t=3.14, p<0.01), but not for muscle ICs (β=-0.01, SD=0.03, t=-0.21, p=0.84) or the brain RV (B<0.01, SD<0.01, t=-1.27, p=0.21). We found no significant interactions between movement intensity and cleaning intensity.

## 4 Discussion

In this study, we investigated the effect of time-domain cleaning and of different levels of mobility on the the outcome of independent component analysis of EEG data. For this, we used eight datasets from six openly available studies and applied the AMICA algorithm using its own automatic sample rejection option with varying strengths. We evaluated the decomposition quality on the basis of the component mutual information, the number of brain and muscle ICs as determined by the ICLabel classifier, as well as the brain IC residual variance in a between-study design. We further evaluated these variables in a within-study design of two of the available datasets. In addition, we measured the SNR in an exemplary application in a within-study design. We hypothesized that increased levels of mobility would lead to decreased decomposition quality, stronger cleaning would lead to an increase in decomposition quality, and higher mobility would require more cleaning to reach the best quality. We found a small, but significant effect of cleaning strength on the decomposition quality. We found no conclusive effect of movement intensity on the decomposition between studies, but the within-study comparison showed a significant effect. We found no evidence that increased mobility requires increased cleaning. Results suggest that differences between studies are stronger than the effect of movement intensity.

### 4.1 Ambiguous effect of movement intensity

Looking at the full between-study comparison, we found no effect of movement intensity in the investigated datasets. We found variations between studies in different metrics but these variations did not seem to depend on the movement intensity when comparing different studies. It could be assumed that higher movement intensities induce more muscle activity to be captured by the AMICA as muscle ICs. Since the number of ICs is limited by the number of channels, an increase in muscle ICs could come at the cost of a lower number of brain ICs. However, this does not seem to be the case in the present study. Neither did the physiological plausibility of the brain ICs vary systematically with movement intensity, as the best and worst scoring datasets were from the same movement intensity group.

This lack of an effect of movement on the ICA decomposition quality could have several reasons: First, it may be the case that there simply is no adverse effect of mobility on data quality. Second, another possible explanation could be that although we attempted to standardize the electrode layouts of the different studies, slight differences in the number and placement of electrodes remained, which could have affected the ICA. It has been shown that the number of electrodes does play a role in the number of resulting brain ICs and the quality of the ICA decomposition (Klug & Gramann, 2021), but we would not expect that effect to be so substantial that it results in large differences like the one observed in the present study. A third possible explanation could be a suboptimal classification of the datasets with respect to movement intensity. We classified the datasets into low, medium, and high movement intensity primarily based on the amount of expected head movement. Future studies should systematically employ alternative movement classification schemas that investigate different kinds of movements and their impact on the recorded EEG data quality. These investigations should also include motion capture to allow an objective criterion of movement intensity. While slow walking might not impact mobile EEG recordings at all, upper torso and arm movements even in seated participants might be associated with head and cap movements that could lead to electrode micromovements associated with non-stationarity of the signal. Thus, even though participants’ mobility is low while sitting and moving their arm, the movement itself might have a stronger impact than walking which can be considered the higher mobility condition. Fourth and last, a major contributing factor to the decomposition quality might not be the mobility of the paradigm itself, but other aspects of the data recording such as the lab environment or the equipment used (Melnik et al., 2017).

The last point is also supported by our post-hoc cluster analysis of the eight available datasets. We found that when attempting to cluster the studies based on their initial data quality (without any cleaning applied), the resultant clusters were not in line with our a priori classification. Rather, studies stemming from the same lab were clustered together: The two *Beamwalking* studies, as well as the two *Spot Rotation* and the *Prediction Error* studies (which were recorded in the same lab) were in the same clusters, respectively. Comparing the AIC scores of the models, we did find that the LMMs that were fitted on the data driven clusters had a significantly better fit than those with our own classification for the dependent variables of mututal information, number of muscle ICs, and residual variance. In sum, we believe that our attempt at a between-study comparison of movement intensity was unsuccessful, and instead within-study comparisons should be done for this investigation to remove uncontrolled variability between labs.

We had the opportunity to test this within-study comparison in addition to our larger comparison across studies since we included two studies that contained both a mobile and a stationary condition, allowing for a direct comparison of the results within the two studies. Indeed, we found a significant effect of mobility in this comparison. Thestudies showed a decrease in the number of brain ICs in the mobile condition and an increase in RV, resulting in a noticeable decrease in the ratio of the number of brain ICs to their mean RV. Although this did not hold true forthe number of muscle ICs, a marginal effect of this reduction in quality was also found in the SNR values of the *Spot Rotation* study.

Taken together, it is likely that in a lab environment where everything is kept constant except the mobility of the participant, an increase in movement intensity will have a negative impact on the data. This negative impact, however, is less pronounced than anticipated, as we did not find it when comparing different studies from different lab setups.

### 4.2 Moderate cleaning improved the decomposition

The impact of cleaning intensity on the quality of the decomposition was significant, but smaller than anticipated. While different datasets scored differently on various metrics, these scores stayed for the most part within the SE range and not all datasets exhibited a noticeable effect. We did observe a positive effect of moderate cleaning on the number of brain ICs and their RV values, but the magnitude was limited, and some datasets exhibited almost no change. Additionally, some datasets required relatively strong cleaning to reach their maxima, while others showed a negative impact of too strong cleaning.

This might be because the AMICA algorithm is more robust than anticipated and suitable for capturing or ignoring artifacts even without substantial cleaning. Especially considering physiological activity not stemming from the brain, removing single samples or small patches from the data does not remove the general activity of these sources. Thus, the AMICA algorithm will have to capture this activity regardless, and removing samples might not help much. Essentially, what researchers may consider artifacts in the data (such as eye movements, muscle activity, or recurrent cable sway from gait) is not necessarily an artifact for the underlying ICA model. If these signals are systematic and can be effectively modeled by the ICA, they will not be removed from the data and neither is it necessary to remove them beforehand. Artifact is a term from the user’s perspective - the model is blind to such labels. Thus, only data that contains large, transient spikes or excessively strong other artifacts that can not easily be modeled by ICA but can be removed in the time-domain would benefit from cleaning. This could for example be time points where the participant was touching the EEG cap or other equipment, or moments where a virtual reality display is taken on or off. However, assuming that these artifacts are limited to periods of breaks or happen before or after the experiment, it might be suitable to just remove all non-experiment segments of the data and perform only minor additional time-domain cleaning before ICA.

If it is essential to capture as many brain components as possible because one is interested in deep or unusual regions of interest and intends to perform source-level analysis, it might be justified to clean the data more strongly. However, in these cases, one must keep in mind that the resulting decomposition will not be able to fully capture the artifacts as it was not computed with them included. This may result in no relevant change in the actual measure to investigate, such as ERPs, as could be seen in the absent effect of time-domain cleaning on the exemplary SNR of the *Spot Rotation* data when all non-brain ICs were removed.

### 4.3 More movement did not require more cleaning

As we expected an adverse effect of movement on the decomposition quality and a positive effect of time-domain cleaning, we also assumed that more cleaning would be necessary for data containing more movement. To our surprise, we found no consistent results indicating an interaction between the movement intensity and the required time-domain cleaning on the resulting decomposition quality. While we did find some indication for main effects, these trends were mostly shared across datasets in direction and magnitude. There were some exceptions but these were single datasets and their trend was usually not shared by the other dataset with the same class of movement intensity, and even if such an effect appeared, it was small. In accordance with our discussion of the expected main effects above, this again suggests that movement and thereby movement artifacts are less impactful on the ICA decomposition than previously assumed, and no substantially different time-domain cleaning is necessary for mobile EEG studies.

### 4.4 Limitations and possible improvements

As a first and major limitation, this study can only discuss the effects of the included datasets and does not necessarily generalize to other lab setups and experimental protocols. It was difficult to find an effect of variations in data processing without controlling for general data quality. A control for data quality, however, is not straightforward and would most likely only be possible when all investigated datasets share the same laboratory setup and recording equipment (Melnik et al., 2017), as well as experiment paradigm (such as an oddball task). Hence, although we tried to find a suitable amount of representative datasets with varying protocols, it would be favorable to have different studies repeat the same protocol in varying movement conditions. A taxonomy of different movement types such as gait, balancing, arm reaching, or tool use would be useful in this case, including a specification of the expected impact of these movement types on the EEG electrodes, ideally confirmed by motion capture. Such a large dataset with consistent recording quality could help shed light on the smaller effects we found, especially since the results contradicted our expectations.

A second limitation is the measure of decomposition quality. We used the number of brain ICs as classified by ICLabel, and their RV as a proxy for decomposition quality, but this approach has two limitations: i) The body of data that ICLabel used to train the classifier did not contain sufficient examples from mobile experiments, meaning that the classification results might not be fully reliable in our context. Extending the classifier to MoBI or mobile EEG studies would alleviate this issue. Such a project is in preparation and the community is invited to contribute, see https://www.icmobi.org. ii) RV values might also be problematic to interpret, especially those of non-brain sources, which is why we did not take those into account. However, especially in the MoBI context, having more physiologically plausible muscle and eye ICs would also be of value, and this is impossible to measure using the current version of dipole fitting in EEGLAB. In the future, this can be done using HArtMuT, a head model that contains sources for eyes and muscles and can thus lead to more reliable estimates of the IC source and its RV (Harmening et al., 2022), but was not yet available at the time of this study. Another option to take into account is to investigate the SNR after data cleaning in more depth. This, however, would also require the same study to be repeated in different movement conditions, akin to the *Spot Rotation* SNR evaluation we performed. This would shed light on more practical implications of the investigated effects. Yet this is still nontrivial to interpret because even in a shared paradigm the SNR of the ERP can change not necessarily due to artifact but due to a change in the neural component, as discussed e.g. in (Ladouce et al., 2019). Hence, although an evaluation based on the SNR of the neural responses would be preferred in principle, it comes with its own interpretational drawbacks and we thus decided to use the ICA-based decomposition quality, but provide the SNR-based information of one within-study-analysis as an addition.

As a corollary to the second limitation, a third limitation is the lack of a ground truth of brain and other physiological and artifactual contributions to the data. Here, a validation study based on properly constructed simulated data would be a valuable tool to investigate the effects in question. However, while working with simulated data is informative, this kind of data is usually cleaner than what researchers find in their real experiments as the complex relationship of movement and muscle activity is hard to simulate accurately. Researchers working with real EEG data must understand the effects of data cleaning and experimental paradigms on this natural data and its ICA decomposition. Although the approach has the shortcoming of an unobtainable ground truth, it is highly valuable to the practical process of data cleaning in many research labs. In a future study, such a comprehensive simulated dataset might be constructed using SEREEGA (Krol et al., 2018), which could then be used to further validate and investigate the questions at hand.

A fourth limitation could be that we only used one method for time-domain cleaning. It is possible that other cleaning options could lead to different results. However, we believe that since the AMICA auto sample rejection uses its own objective metric, it is unlikely that the cleaning results will be substantially improved when using other algorithms. A separate investigation of the effect of a proposed time-domain cleaning algorithm in comparison with the AMICA auto sample rejection found no noticeable difference (Klug et al., 2022).

### 4.5 Conclusions

In our investigation of the effect of time-domain cleaning and movement intensity on the quality of the ICA decomposition, we found evidence that cleaning has a moderately positive effect on the decomposition quality, and movement has a negative effect on the decomposition quality when compared within studies. The effect of movement was not found between studies, pointing to the fact that lab setups, equipment, and possibly the paradigm itself might have a greater impact on the decomposition quality than the movement intensity. Additionally, while we did find evidence that moderate cleaning prior to ICA computation improves the decomposition, this effect was far weaker than anticipated and it did not vary systematically with movement intensity in our study. This suggests that the AMICA algorithm is very robust and can handle artifacts even with limited data cleaning.

We thus recommend not to remove substantial parts of the data using time-domain cleaning before running AMICA. Moderate amounts of cleaning such as 5 to 10 iterations of the AMICA sample rejection starting after 2 iterations with the default 3 SDs as threshold and 3 iterations between rejections will likely improve the decomposition in most datasets, irrespective of the movement intensity. This results in around 5% removed data. Only in special circumstances, strong cleaning will be relevant and more beneficial.

## Acknowledgments

We are thankful to the researchers who made their datasets freely available. Without them, this investigation would not have been possible.

## Data availability statement

The data used in this study is available for download as laid out in the *Datasets* section.

